# Single-cell lineage trajectory defines CDK inhibitor-sensitive cells-of-origin in esophageal squamous cell cancer

**DOI:** 10.1101/2025.10.11.681802

**Authors:** Kyung-Pil Ko, Jie Zhang, Sohee Jun, Jae-Il Park

## Abstract

Understanding the cells of origin is essential for overcoming therapy resistance in esophageal squamous cell carcinoma (ESCC). We utilized machine learning-based single-cell trajectory analysis on 4NQO-induced murine models and genetically engineered organoids to identify multiple distinct cell clusters that serve as cellular origins of ESCC. Gene regulatory network analysis of these populations indicated activation of stem/progenitor cell regulators, including PRRX2 and CEBPβ. Translating these findings, a transcriptome-based drug repurposing screen identified five chemical candidates, four of which are potent Cyclin-Dependent Kinase (CDK) inhibitors, aligning with the frequent loss-of-function mutations in *TP53* and *CDKN2A* observed in ESCC. Notably, CDK inhibitors markedly inhibit ESCC cell proliferation. This research delineates the potential cellular origins of ESCC and their key regulons, thereby pioneering a single-cell-derived therapeutic strategy that exposes vulnerabilities in tumor-initiating cells.

## Introduction

Esophageal cancer is divided into esophageal squamous cell cancer (ESCC) and esophageal adenocarcinoma (EAC). ESCC accounts for over 80% of all cases of esophageal cancer and has a poor prognosis because of a lack of symptoms in the early stages ^1^. The overall five-year survival of patients with esophageal cancer ranges from 10% to 25% ^2^. ESCC develops from squamous dysplasia as an invasive, typical histologic precursor lesion ^3^, which impedes the early detection of the lesion and consequently results in late diagnosis, thereby adversely affecting patient survival.

Given that early diagnosis of ESCC may bring better outcomes ^2^, understanding the genetic, cellular, and molecular mechanisms of esophageal neoplasia and ESCC initiation is imperative, which may improve the detection, diagnosis, treatment, and prevention of ESCC. However, the biology of ESCC initiation remains elusive.

The esophageal epithelium is composed of a proliferative basal layer and differentiated suprabasal layers of epithelial cells ^4^. The basal epithelium of the murine esophagus contains both proliferating stem and transit-amplifying cells that self-renew and differentiate over the tissue’s lifespan ^5^. Three-dimensional organoids, which simulate physiological and pathological organ processes, have become a promising tool for studying stem cells and diseases ^6,7^. We recently re-formulated the culture media for esophageal organoids (EOs), which are cost-effective and superior to conventional ones, and established a new EO system that mimics ESCC’s early lesion ^8,9^. To understand the genetic mechanism of ESCC initiation, we genetically engineered EOs and identified the key genetic determinants (loss-of-function of *TP53, CDKN2A*, and *NOTCH1*) that initiate ESCC tumorigenesis and immune evasion ^10^.

ESCC is mainly treated by surgery, while radiotherapy, chemotherapy, and chemoradiotherapy have limited efficacy ^11,12^. Since cells-of-origin of cancer (also called cancer stem and progenitor cells) are likely responsible for therapy resistance, relapse, and metastasis ^13,14^, targeting cells-of-origin has been highlighted to overcome the limitations in cancer treatment ^15^. To this end, many studies have focused on identifying specific biomarkers for such cellular origins ^16,17^. Nonetheless, the current understanding of the cellular origin of cancer (especially in solid tumors) is still insufficient to pinpoint the specific cells for targeted treatment. This is partly due to the limitation of current approaches, which heavily depend on a single or a few biomarkers to define self-renewing tumor cells.

To overcome this, we employed single-cell transcriptomics and identified the earliest cell clusters serving as potential cellular origins of ESCC tumorigenesis. By leveraging genetically engineered organoids and single-cell transcriptomics, this study aimed to determine whether pharmacological interventions targeting cellular origins, as defined by single-cell transcriptomics, are sufficient to suppress ESCC tumorigenesis, laying a new foundation for developing ESCC therapies.

## Results

### Identification of multiple cells-of-origin in the ESCC models

To pinpoint the cells of origin responsible for ESCC tumorigenesis, we employed single-cell transcriptomics-based approaches. We analyzed single-cell RNA sequencing (scRNA-seq) datasets derived from the esophageal tissues of mice treated with 4NQO (4-nitroquinoline 1-oxide), a potent carcinogenic agent ^18^. This 4NQO-treated mouse model exhibits a progressive series of lesions that faithfully mimic the developmental stages of ESCC tumorigenesis, encompassing inflammation, hyperplasia, dysplasia, and cancer in situ (CIS) (Fig. 1A). Notably, our recent studies have demonstrated a high degree of similarity between the single-cell transcriptomes of 4NQO-treated ESCC mouse models and those of human ESCC ^10,19^. Transcriptomes of cells from five different stages were integrated and annotated for isolating epithelial cells from other cell types, such as fibroblasts (Fig. 1B). Cell clusters that are dominant in the normal dataset were annotated as normal epithelial cells (Epi 1–8), while CIS-abundant cell clusters were named as neoplastic cells (Neo 1–11).

**Figure 1.**
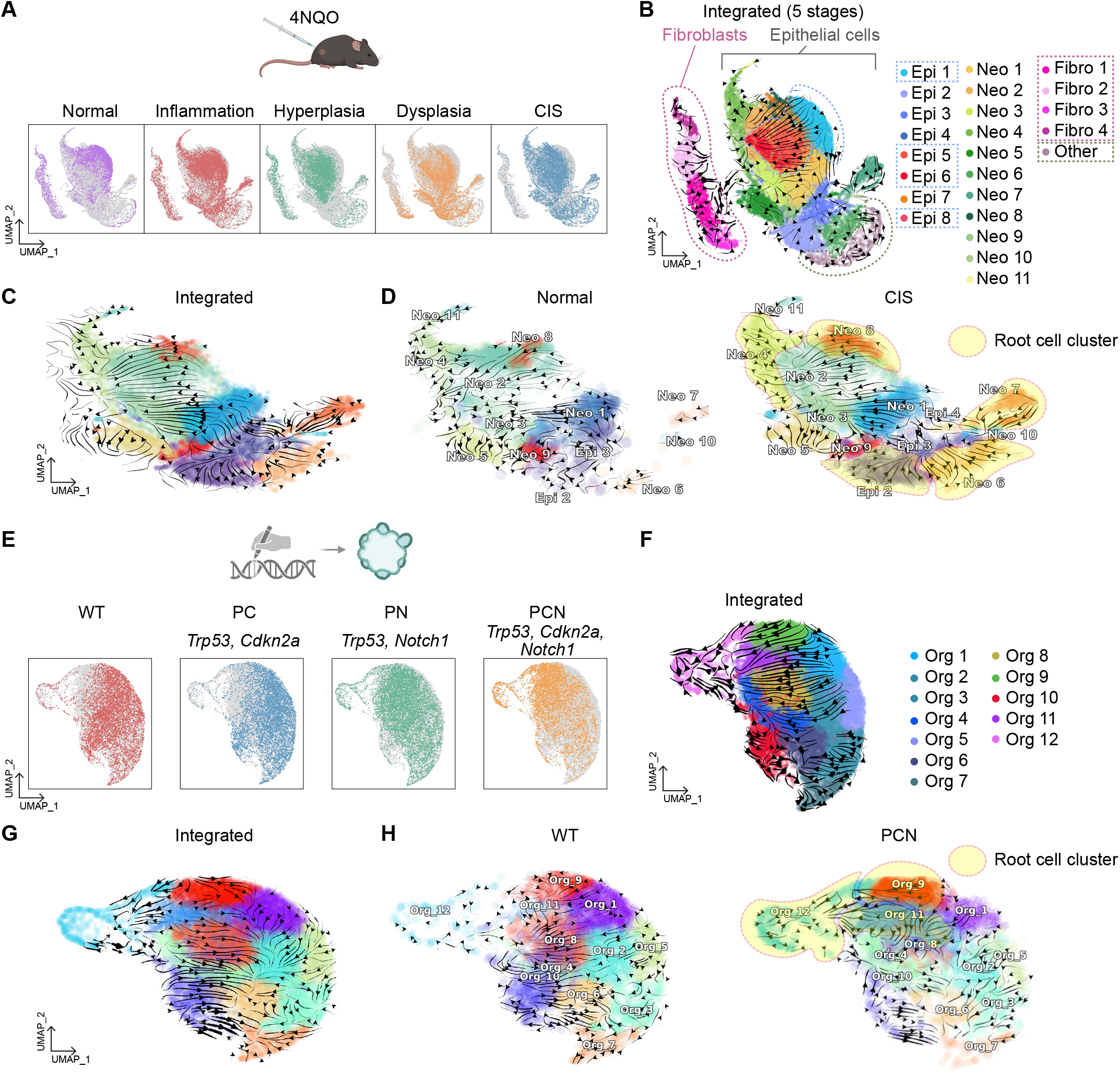
Identification of cells-of-origin clusters of ESCC. **A**. UMAPs from the integrated dataset from 4NQO-treated mouse esophageal samples. CD45(-) cells from Normal, Inflammation, Hyperplasia, Dysplasia, and CIS datasets were integrated using the Harmony package. **B**. RNA velocity-based UMAP from the integrated dataset from 4NQO-treated mouse esophageal samples (Normal, Inflammation, Hyperplasia, Dysplasia, and CIS). RNA velocity was calculated with the scVelo package. Proliferating cells or differentiated cells were annotated with Epi 1–8, and the fibroblasts were annotated with Fibro 1-4 based on the marker gene expression. Cell clusters that have mixed gene expression patterns of proliferating cells and stratified cells were annotated with Neo 1-11, and a cluster that does not express epithelial cell markers was named Other. **C**. Normal cell lineage-related (Epi 1, 5∼8), Fibroblast (Fibro 1∼4), and Other clusters were excluded, and the RNA velocity was calculated again using the Dynamo package. **D**. Normal and CIS datasets were separated from the integrated data, and the cell differentiation streams were displayed using Dynamo. Cells-of-origin clusters (Epi 2, Neo 4, 6, 7, and 8) specific to CIS were highlighted. **E**. The Integrated dataset was segregated with each genotype (PC, PN, and PCN) and shown on the UMAP. **F**. The UMAP, showing the integrated organoid datasets (WT, PC, PN, and PCN), and RNA velocity calculated by scVelo, was displayed on the UMAP. **G**. RNA velocity was recalculated with Dynamo. **H**. RNA velocity of WT and PCN subsets from the integrated dataset was displayed separately. Cells-of-origin clusters (Org_9, 11, and 12) enriched in PCN were highlighted.

Next, we harnessed the power of cell lineage trajectory inference analysis, a cutting-edge method, on the scRNA-seq datasets sourced from 4NQO ESCC mouse models. Employing three distinct analytical packages, namely RNA velocity (scVelo; based on RNA splicing), CytoTRACE (based on RNA content), and Dynamo (utilizing machine learning) ^20-22^, we visualized cell lineage trajectories (Fig. 1B). To directly compare the change of cellular trajectories in normal and CIS esophagi, we isolated only Normal and CIS datasets (Fig. 1C). In contrast to the cells of origin identified in normal esophagi, the CIS stage displayed five distinctive cells of origin clusters, Neo 4, 6, 7, 8, and epi 2 (Fig. 1C, D).

Moreover, we extended our analysis to include scRNA-seq datasets from genetically engineered EOs that we recently pioneered ^10^. These EOs encompass normal EOs (wild-type [WT]), PC (*Trp53 Cdkn2a* knockout [KO]), PN (*Trp53 Notch1* KO), and PCN (*Trp53 Cdkn2a Notch1* triple KO) EOs, with the latter demonstrating the development of invasive ESCC tumors ^10^ (Fig. 1E).

We integrated four different organoid datasets and annotated each epithelial cell cluster (Org 1–12) (Fig. 1F). In this integrated dataset, two cell clusters, Org 2 and Org 9, were identified as the major cells of origin. To exclude masking effect from PC and PN data, we isolated only WT and PCN datasets and compared their cell lineages (Fig. 1G). Intriguingly, the WT organoid exhibited an opposite cellular lineage direction compared to the previous integrated dataset. The Org 4 cell cluster was the cell-of-origin of normal esophageal organoids and differentiates into others (Fig. 1H). Remarkably, three cell clusters, Org 9, 11, and 12, were revealed as initiating cell clusters of PCN EOs, confirming multiple cellular origins in the neoplastic cells (Fig. 1H). The lineage directions of PCN were similar to those of the previous integrated dataset, likely reflecting the transcriptomic proximity of PC and PN to the PCN data. These results indicated that ESCC cells are heterogeneous, likely driven by various cells of origin.

### Gene regulatory networks of cells of origin

Next, we analyzed the gene regulatory networks characterizing each cell-of-origin cluster within the CIS scRNA-seq dataset by using the pySCENIC package ^23^. Among the top ten regulons identified following five rounds of pySCENIC analyses for the cells-of-origin clusters (Epi2, Neo4, 6-8), we pinpointed eight regulons (Prrx2, Cebpb, Zbtb7b, Snai2, Tfap2a, Tfap2c, Trp63, and Snai3) that were consistently activated in the cells-of-origin clusters of CIS, but not in the normal dataset (Fig. 2A, B). PRRX2 serves as a marker for pituitary stem/progenitor cells ^24,25^, while CEBPβ plays a regulatory role in hematopoietic and skeletal stem cells ^26,27^. SNAI2 is known for its control over epidermal progenitor cells ^28^, and TFAP2A/C is associated with the regulation of pluripotent stem cell differentiation ^29^. Moreover, p63/TP63 is recognized for its role in modulating the proliferation of epithelial stem and progenitor cells ^30^.

**Figure 2.**
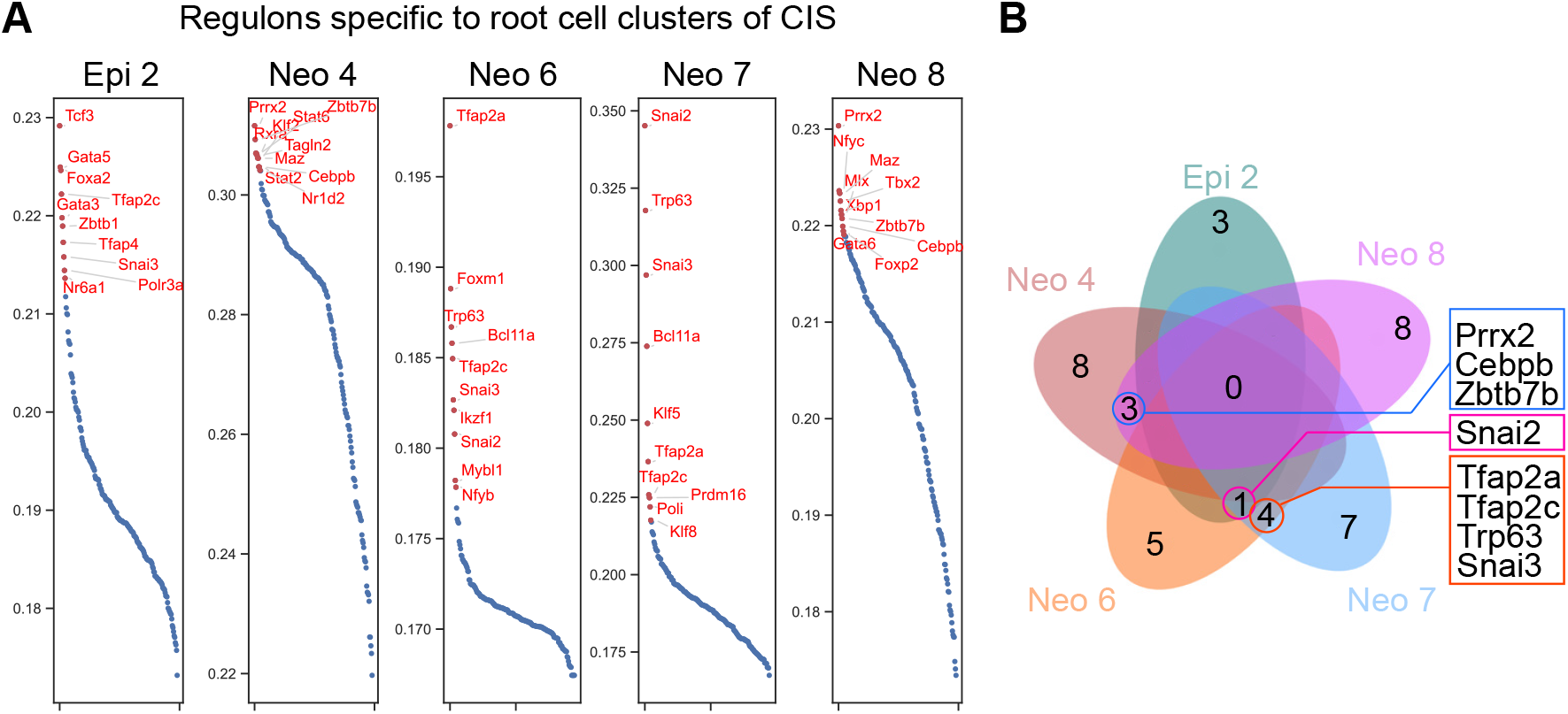
Gene regulatory network specific to cells-of-origin clusters of ESCC. **A**. The gene regulatory network (GRN) was calculated from the cells-of-origin clusters of the CIS dataset. The pySCENIC package was used for regulon analysis, and the top 10 regulons for each cluster were displayed based on their regulon scores. Each cluster was calculated 5 times with pySCENIC. **B**. The top 10 regulons from 5 rounds of analysis were compared to identify the shared regulons within each cluster. Only shared regulons from 5 rounds of each cluster were compared with those of other cells-of-origin clusters, and eight regulons (Prrx2, Cebpb, Zbtb7b, Snai2, Tfap2a, Tfap2c, Trp63, and Snai3) were identified to overlap in at least two clusters.

### Signaling pathways in the ESCC initiating cell clusters

Since a significant number of epithelial cells from the normal esophagus were also observed in the CIS-enriched initiating cell clusters, such as Neo 4 and Neo 8, we compared the signaling pathways between the normal and CIS datasets within these cell clusters. In the Neo 4 cell cluster, cytokine-mediated pathways, including IL-17 and TNF signaling, were enriched in the CIS compared to normal esophagus, indicating more active cell-cell interactions in CIS than normal cells (Fig. 3A). In the Neo 8 cluster, mitochondrial electron transport and oxidative phosphorylation pathways were stronger in CIS than in normal cells, suggesting enhanced energy production through OXPHOS metabolism. These results suggest that ESCC-initiating cells are heterogeneous, with distinct inter- and intracellular processes.

**Figure 3.**
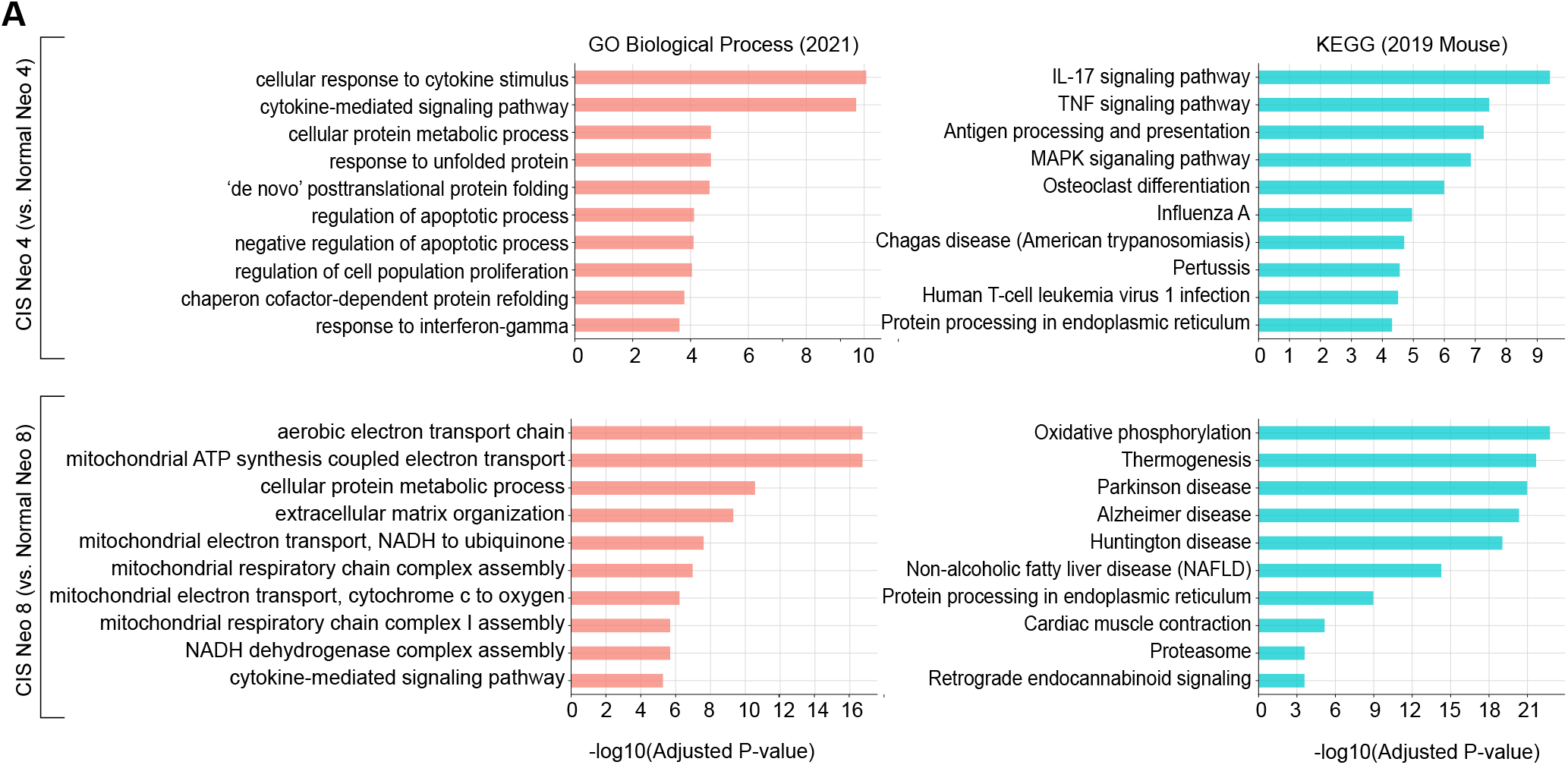
GSEA analysis of initiating cells between normal and CIS. **A-B**. GSEA from differentially expressed genes (DEGs) between CIS and Normal cells within the Neo 4 (A) and Neo 8 (B) clusters. The top 10 significant pathways from GO Biological Process and KEGG databases are shown.

### Identification of drug candidates potentially targeting cells-of-origin of ESCC

Having identified heterogeneous cellular processes in the initiating cell clusters through scRNA-seq analysis (Fig. 3), our subsequent objective was to pinpoint chemicals or drugs that specifically, yet integratively, target these cell-of-origin clusters. To achieve this, we employed a transcriptome-based drug repurposing algorithm, as recently performed ^31,32^. In a manner akin to the Connectivity Map (CMap) ^33^, we conducted a data-driven analysis focusing on drug mode-of-action and drug repositioning. Rather than considering the entire transcriptome, we generated lists of differentially expressed genes (DEGs) that were unique to each of the cells-of-origin clusters. Subsequently, we utilized the L1000CDS^2^ tool ^34^ and identified five promising chemical candidates (CGP60474, Flavopiridol, AZD-5438, SNS-032, and daunorubicin) with their potential to target the cells-of-origin clusters (Fig. 4A). Of note, CGP60474 is an inhibitor of cyclin-dependent kinases (CDK) and PKC. Flavopiridol, also known as alvocidib, holds FDA approval as a CDK inhibitor. AZD-5438 similarly acts as a CDK inhibitor, blocking CDK1, CDK2, and CDK9. Meanwhile, SNS-032 is recognized for its inhibition of CDK2, CDK7, and CDK9. Daunorubicin, an FDA-approved agent, functions as a DNA intercalating agent akin to doxorubicin, primarily used in the treatment of leukemia. Among these five chemicals, CGP-60474 (with an IC_50_ of 0.011 and 0.078 mM) and Flavopiridol (with an IC_50_ of 0.066 and 0.318 mM) exhibited noteworthy growth inhibitory effects on TE-1 and TE-12 human ESCC cell lines (Fig. 4B-D).

**Figure 4.**
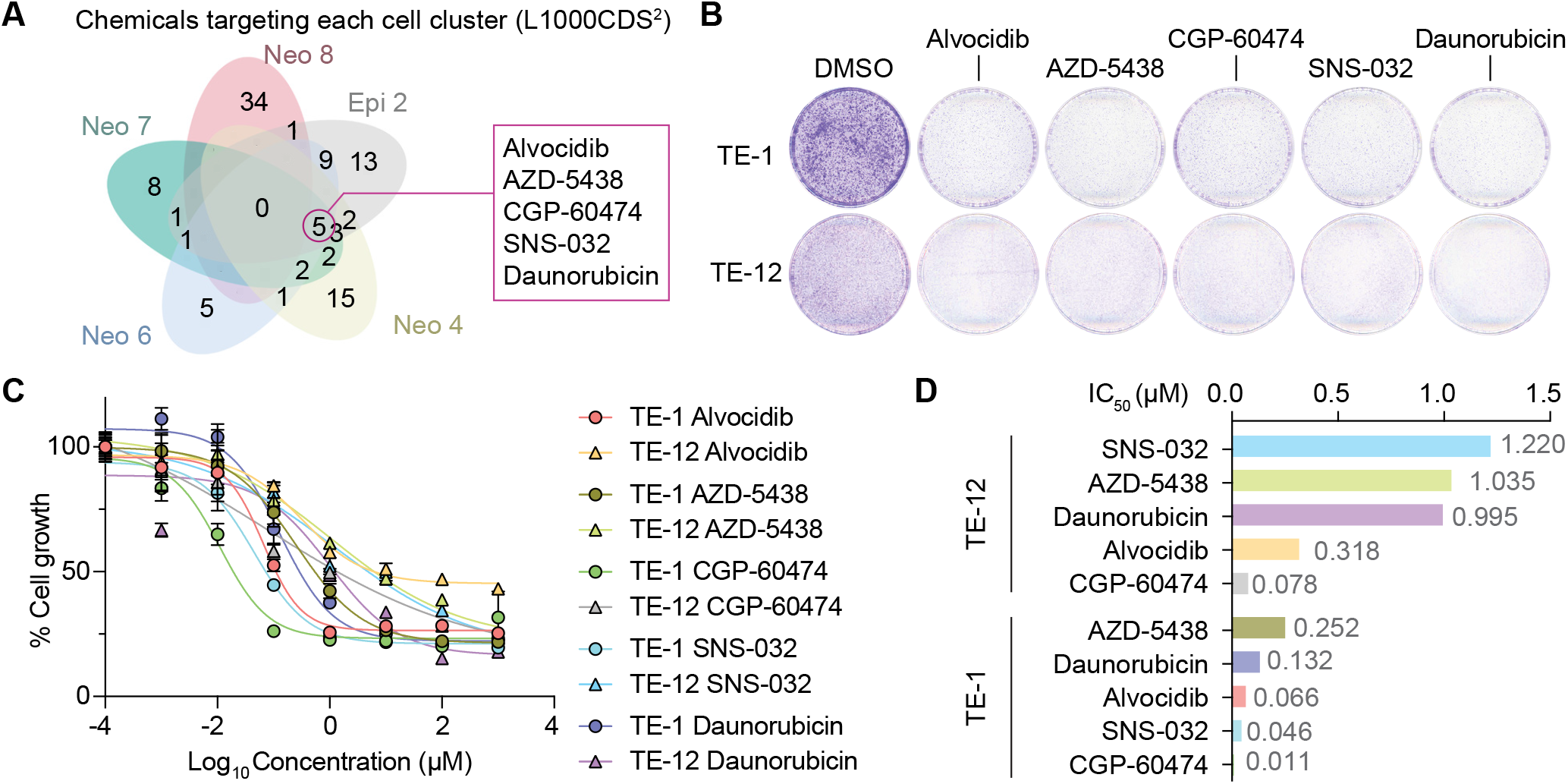
Identification of chemicals targeting cells-of-origin of ESCC. **A**. Drug candidates were identified from L1000CDS^2^ analysis. DEGs from CIS and Normal in each cell-of-origin cluster were used for analysis, and five chemicals were shared in four clusters. **B**. Colony formation assay showing the growth inhibitory effect of five drugs in TE-1 and TE-12 human ESCC cell lines. Cells were stained with Crystal violet 48 h after chemical/drug treatment (10 μM). Representative images are shown (n = 3). **C**. The impact of five drugs/chemicals (alvocidib, AZD-5438, CGP-60474, SNS-032, and daunorubicin) on the cell growth of TE-1 and TE-12 cells was analyzed using the CCK-8 assay. Different doses (10^-4^ μM – 10^3^ μM) were tested after 48 h of treatment. DMSO (vehicle) served as control. **D**. IC_50_ values of the candidate drugs are shown.

## Discussion

Understanding the cells of origin has mainly been emphasized in developmental biology and regenerative medicine. Starting from long-term labeled cell detection, by using genetically engineered mouse models, genetic labeling-based cell lineage tracking and cell ablation approaches have revealed several tissue stem and progenitor cells ^35-41^.

The concept of targeting cells of origin, especially cancer stem cells, has emerged as a promising strategy to combat therapy resistance, reduce relapse rates, and impede metastasis in various malignancies ^13,14,42-44^. Nevertheless, the current understanding of cancer cell origins, particularly in solid tumors, remains incomplete, hindering the precise identification and targeting of these critical cellular populations. This limitation partly stems from the conventional methodologies, which predominantly rely on a single or a limited set of biomarkers for the identification and characterization of self-renewing tumor cells. The recent advent of single-cell transcriptomics and genomics has enabled a comprehensive understanding of cellular classifications, lineage specification, and plasticity in development, tissue regeneration, and tumorigenesis ^8,10,19-22,32,45-49^.

To identify the cells of origin in ESCC, we analyzed scRNA-seq datasets of 4NQO ESCC mouse models ^18^ and genetically engineered organoids. A 4NQO ESCC mouse model recapitulates ESCC tumorigenesis, including inflammation, hyperplasia, dysplasia, and CIS, as analyzed by scRNA-seq datasets ^10^. By using scVelo, CytoTRACE, and Dynamo, we identified the distinct cell lineage trajectories of CIS compared to those of normal esophagus. Machine learning-based Dynamo best identified the five cell clusters (Neo 2, 4, 6, 8, and epi 2) serving as cells-of-origin in CIS (Fig. 1). Further analyses identified the key gene regulatory networks specifically activated in the cells-of-origin of CIS compared to the normal esophagus (Fig. 2, 3). Notably, we identified five drug or chemical candidates (CGP60474, Flavopiridol, AZD-5438, SNS-032, and daunorubicin) that significantly inhibited tumor cell growth (Fig. 4).

Interestingly, four of these chemical candidates (CGP60474, Flavopiridol, AZD-5438, and SNS-032) were known to be potent and selective inhibitors of CDKs, implying therapeutic vulnerabilities in cyclin-dependent cell cycle regulation. This might be originated from clonal evolution driven by consensus genetic alterations during ESCC development. Unlike other types of cancer, ESCC patients show an extremely high frequency of *TP53* mutation and loss of *CDKN2A* genes, which are more than 90% and 70%, respectively.^10,50-52^ Considering their overlapping roles in RB1-mediated cell cycle control, better therapeutic outcomes for ESCC can be achieved by targeting cyclin-dependent cell cycle. Consistent with our findings, more than 62% of ESCC cases exhibited genetic profiles of disrupted G1/S transition control ^53^.

There were several preclinical trials to suppress ESCC growth using CDK inhibitors alone or combinatorial therapy ^54-56^. ESCC cell proliferation was significantly reduced by dalpiciclib and SNS-032, and sensitivity to radiotherapy or chemotherapy was enhanced by CDK inhibitors, such as palbociclib. Although CDK inhibitors show promising results in ESCC cell lines, further research is necessary to determine whether their tumor suppressive effects stem from blocking cancer cell stemness or from cytostatic mechanisms, using patient-derived xenografts or organoids.

Together, this study elucidates the potential cellular origins and driving regulons involved in ESCC tumorigenesis. Notably, we have translated these findings into a therapeutic strategy by pioneering the application of single-cell transcriptome data to identify compounds that could suppress cancer cell stemness, thereby potentially targeting the cells-of-origin of tumor heterogeneity, metastatic progression, and therapeutic resistance. This study provides a new foundation for advancing the understanding and treatment of ESCC.

## Author contributions

K.-P.K.: Conceptualization, methodology, investigation, software, analysis, data curation, writing, visualization, funding acquisition; J.Z.: Investigation; S.J.: Investigation; J.-I.P.: Conceptualization, analysis, writing, visualization, supervision, project administration, funding acquisition

## Acknowledgments

This work was supported by the National Cancer Institute (CA286761 to K.-P.K. and CA278971 to J.-I.P.). We thank Dr. Shumei Song for providing us with the TE-1 and TE-12 cell lines.

## Methods

### Cell culture

Human ESCC cell lines TE-1 and TE-12 were kindly provided by Dr. Shumei Song. Cells were maintained in RPMI 1640 supplemented with 10% fetal bovine serum (FBS) at 37°C under 5% CO2 atmosphere. New vials of frozen stocks were routinely thawed and cultured for experiments, minimizing long-term passaging effects.

### Colony formation and cell viability assays

For colony formation assays, 1 × 10□ cells were plated in 60-mm dishes and cultured with medium refreshed every two days. Colonies were fixed with methanol for 20 minutes, rinsed with distilled water, and stained using 0.05% crystal violet. After three additional washes, plates were air-dried and imaged for quantification. For cell viability assays, 1 × 10^3^ cells/well were seeded in 96-well plates. After 24 hours, the medium was replaced with drug-containing medium at varying concentrations, followed by 48 hours of incubation. Cell viability was assessed using the CCK-8 reagent (Dojindo) with a 4-hour incubation, and absorbance was measured at 450 nm (reference: 630 nm). Each condition was tested in triplicate.

### Preprocessing scRNA-seq data

Mouse 4NQO-induced ESCC scRNA-seq data were downloaded from the Genome Sequence Archive in BIG Data Center (Beijing Institute of Genomics, Chinese Academy of Sciences, http://gsa.big.ac.cn) under the accession number CRA002118^18^. Organoid datasets were generated in our prior study ^10^. Data preprocessing was conducted using Scanpy (version 1.10.4)^57^. Quality-control filters excluded cells expressing fewer than 100 genes and genes detected in fewer than three cells.

### RNA Velocity–Based Lineage Trajectory Inference

RNA velocity, latent time, and PAGA-directed trajectories were analyzed using scVelo^58^. Briefly, spliced and unspliced Cell Ranger-generated matrices were merged, and genes with fewer than 20 counts were filtered. The top 2,000 highly variable genes were selected. Data were size-normalized, log-transformed, and neighborhood moments computed. RNA velocities were estimated using the dynamical model and visualized on the UMAP embedding. For Dynamo analysis, The velocity vector field was constructed and characterized for topological features including curl, divergence, acceleration, and curvature using Dynamo package^22^.

### Regulon Analysis and Visualization

Transcriptional regulatory network inference was performed using pySCENIC^59^ to infer transcription factor-target interactions and assess regulon activity at the single-cell level. to predict transcription factor–target interactions and evaluate regulon activity at the single-cell level. AUCell scores were computed per regulon and cell cluster to derive regulon specificity scores. To confirm consistency, analyses were repeated five times for each cluster, and the most prominent regulons were identified for visualization.

### Gene Set Enrichment Analysis

Gene set enrichment analysis was carried out using the fgsea package^60^ in R to identify significantly enriched biological processes. Genes from differential-expression analyses were ranked by scores obtained from Scanpy’s rank_genes_groups function and input as pre-ranked lists. msigdbr was used to access hallmark and curated gene sets (databases: GOBP and KEGG).

### Code availability

The code used to reproduce the analyses described in this manuscript can be accessed via GitHub

(https://github.com/jaeilparklab/ESCC_CDKi).

